# Computational metabolomics hints at the relevance of glutamine metabolism in breast cancer

**DOI:** 10.1101/370221

**Authors:** Lucía Trilla-Fuertes, Angelo Gámez-Pozo, Elena López-Camacho, Guillermo Prado-Vázquez, Andrea Zapater-Moros, Rocío López-Vacas, Jorge M Arevalillo, Mariana Díaz-Almirón, Hilario Navarro, Paloma Maín, Enrique Espinosa, Pilar Zamora, Juan Ángel Fresno Vara

## Abstract

Metabolomics has a great potential in the development of new biomarkers in cancer. In this study, metabolomics and gene expression data from breast cancer tumor samples were analyzed, using (1) probabilistic graphical models to define associations using quantitative data without other *a priori* information; and (2) Flux Balance Analysis and flux activities to characterize differences in metabolic pathways. On the one hand, both analyses highlighted the importance of glutamine in breast cancer. Moreover, cell experiments showed that treating breast cancer cells with drugs targeting glutamine metabolism significantly affects cell viability. On the other hand, these computational methods suggested some hypotheses and have demonstrated their utility in the analysis of metabolomics data and in associating metabolomics with patient’s clinical outcome.

## Introduction

Breast cancer is one of the most common malignancies, with 266,120 estimated new cases and 40,920 estimated deaths in the United States in 2018 (Siegel et al, 2018). In clinical practice, the expression of hormonal receptors and HER2 allows the classification of this disease into three groups: hormonal receptor-positive (ER+), HER2+ and triple negative (TNBC).

Metabolomics, a technique focused in the holistic study of the metabolites present in a biological system, is considered the most recent-omics. It consists of measuring the entire set of metabolites present in a biological sample (Fiehn, 2002). The most common techniques in metabolomics experiments are mass spectrometry-related methods, which are based on the mass/charge relationships of each metabolite or its fragments (Emwas, 2015). Recent advances in this technique allow the measurement of thousands of metabolites from minimal amounts of biological samples (Emwas, 2015; Fuhrer & Zamboni, 2015). Therefore, metabolomics is a promising tool for the development of new biomarkers (Gowda et al, 2008).

We used two different methods to merge metabolomics and gene expression data in breast cancer. In previous studies, we used probabilistic graphical models (PGMs) to study differences between breast tumor subtypes and to characterize muscle-invasive bladder cancer at a functional level using proteomics data (Gámez-Pozo et al, 2015; Gámez-Pozo et al, 2017; Sánchez-Navarro et al, 2010). Flux Balance Analysis (FBA), however, is a method that has been widely used to study biochemical networks (Varma & Palsson, 1995). FBA predicts the growth rate or the rate of production of a given metabolite (Orth et al, 2010), and it has previously been used to characterize breast cancer cell responses against drugs targeting metabolism (Trilla-Fuertes et al, 2018). In this study, flux activities were proposed as a feasible method to compare flux patterns in metabolic pathways.

Glutamine has a relevant role in tumor metabolism. The entrance of glutamine in the tricarboxylic acid cycle (TCA) generates lactate, a process known as glutaminolysis. The metabolism of glutamine serves to maintain the availability of non-essential aminoacids and to maintain TCA intermediates while NADH is generating (DeBerardinis et al, 2007). Glutamine is necessary to cellular proliferation and its metabolism is regulated by the levels of *MYC* oncogene (Eagle et al, 1956; Wise et al, 2008).

In the present study, metabolomics and gene expression data from 67 fresh tissue samples (Terunuma et al, 2014) were analyzed through PGMs and FBA. Our aim was to find associations between metabolomics and gene expression data and the characterization of breast cancer from a metabolomics point of view.

## Results

### Patient characteristics

With the aim of study the relationships between metabolomics, gene expression, and FBA results, metabolomics and gene expression data, analyzed by mass-spectrometry and microarrays GeneChip Human Gene 1.0 ST (Affymetrix) respectively and published by Terunuma et al., were analyzed (Terunuma et al, 2014).

A total of 67 tumor fresh tissue samples from patients with breast cancer were studied. This patient cohort comprises 33 ER+ and 34 ER- (of which 14 were also TNBC). The median follow-up was 50 months, and 31 deaths occurred during this time. No significant differences regarding overall survival (OS) were observed between patients with ER+ or ER-tumors. Patient characteristics are shown in Table 1.

**Table 1:** Patient characteristics.

|  | n (%) | ER+ | ER- |
| --- | --- | --- | --- |
| **Number of patients** | 67 | 33 | 34 |
| **Age (years)** |  |  |  |
| Median | 51 | 57 | 48 |
| Range | 30–93 | 34–93 | 30–75 |
| **TNM stage** |  |  |  |
| I | 6 (9%) | 4 (12%) | 2 (6%) |
| II | 2 (3%) | 1 (3%) | 1 (3%) |
| IIA | 23 (35%) | 12 (37%) | 11 (32%) |
| IIB | 21 (31%) | 7 (21%) | 14 (41%) |
| IIIA | 9 (13%) | 5 (15%) | 4 (12%) |
| IIIB | 6 (9%) | 4 (12%) | 2 (6%) |
| **N category** |  |  |  |
| pN0 | 37 (55%) | 17 (52%) | 20 (59%) |
| pN1 | 24 (35%) | 13 (39%) | 11 (32%) |
| pN2 | 5 (8%) | 3 (9%) | 2 (6%) |
| Missing | 1 (2%) | 0 (0%) | 1 (3%) |
| **Grade** |  |  |  |
| G1 | 8 (12%) | 8 (24%) | 0 (0%) |
| G2 | 20 (30%) | 14 (43%) | 6 (18%) |
| G3 | 29 (43%) | 7 (21%) | 22 (64%) |
| Missing | 10 (15%) | 4 (12%) | 6 (18%) |
| **Neoadjuvant therapy** |  |  |  |
| Yes | 6 (9%) | 2 (6%) | 4 (12%) |
| No | 50 (75%) | 26 (79%) | 24 (70%) |
| Missing | 11 (16%) | 5 (15%) | 6 (18%) |

### Analysis of metabolomics data

After Kaplan-Meier analysis, 29 metabolites were found related to OS (p<0.05) (Sup Table 1).

Then, an OS predictor using this metabolomics data was built. This metabolite-based signature included five metabolites: glutamine, 2-hydroxypalmitate, deoxycarnitine, butyrylcarnitine and glycerophosphorylcholine (p-value =0.003, hazard ratio [HR] = 0.34, 50:50%) (Fig 1). A multivariate analysis showed that the predictor provided additional prognostic information to clinical data (S1 Table).

**Fig 1:**
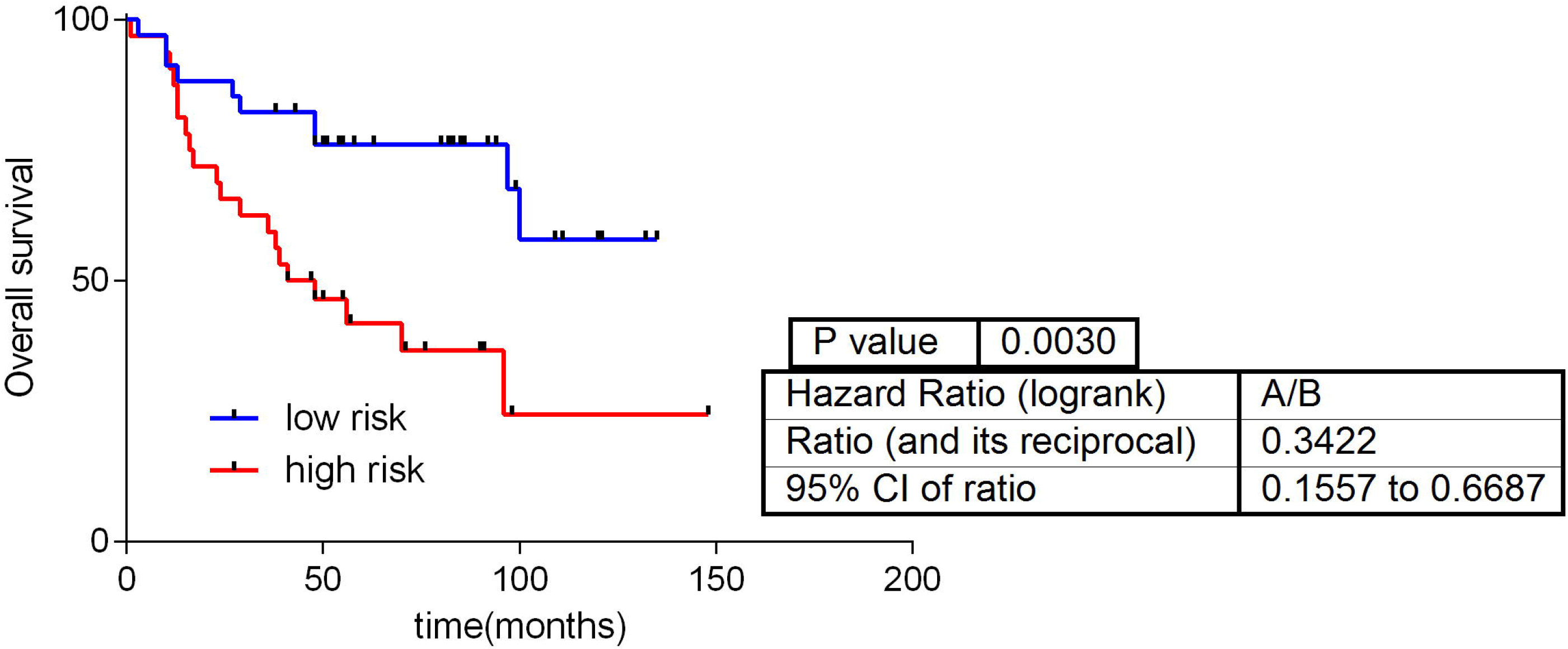
Predictive signature built using metabolomics data.

Metabolomics data without using any a priori information were analyzed through PGM. Metabolomics database, including information about 536 metabolites, was reduced to 237 metabolites due to quality criteria. A main metabolic pathway was assigned to each functional node of the resulting network using IMPaLA. IMPaLA is a tool that allows ontology analyses based on metabolic pathways instead of genes. Strikingly, this network had a functional structure, grouping the metabolites into metabolic pathways as it has been previously shown for gene and protein PGMs (de Velasco et al, 2017; Gámez-Pozo et al, 2015; Gámez-Pozo et al, 2017; Trilla-Fuertes et al, 2018). Five functional nodes were defined, each with a different overrepresented metabolic pathway (Fig 2).

**Fig 2:**
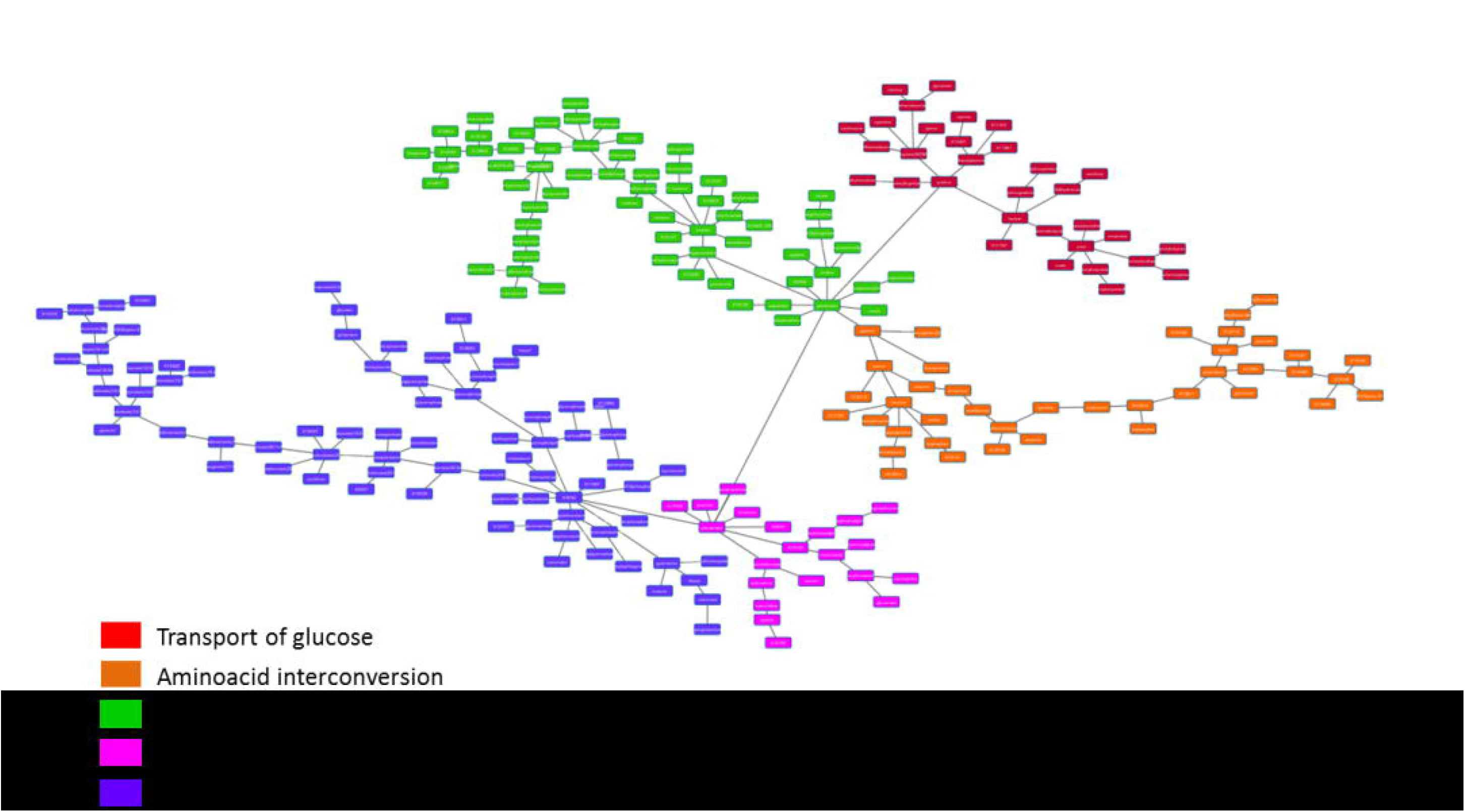
Probabilistic graphical model from metabolomics data.

The activity of each functional node was calculated as previously described and comparisons between ER+ and ER-were done (de Velasco et al, 2017; Gámez-Pozo et al, 2015; Gámez-Pozo et al, 2017; Trilla-Fuertes et al, 2018). Significant differences were found between ER+ and ER-tumors regarding lipid and purine metabolism (p<0.05) (S1 Fig).

Moreover, the lipid metabolism functional node showed prognostic value in this cohort (p =0.045, HR = 0.48, 50:50%) (Fig 3). However, a multivariate analysis showed that the predictor do not add additional prognostic information to that provided by clinical features (S2 Table).

**Fig 3:**
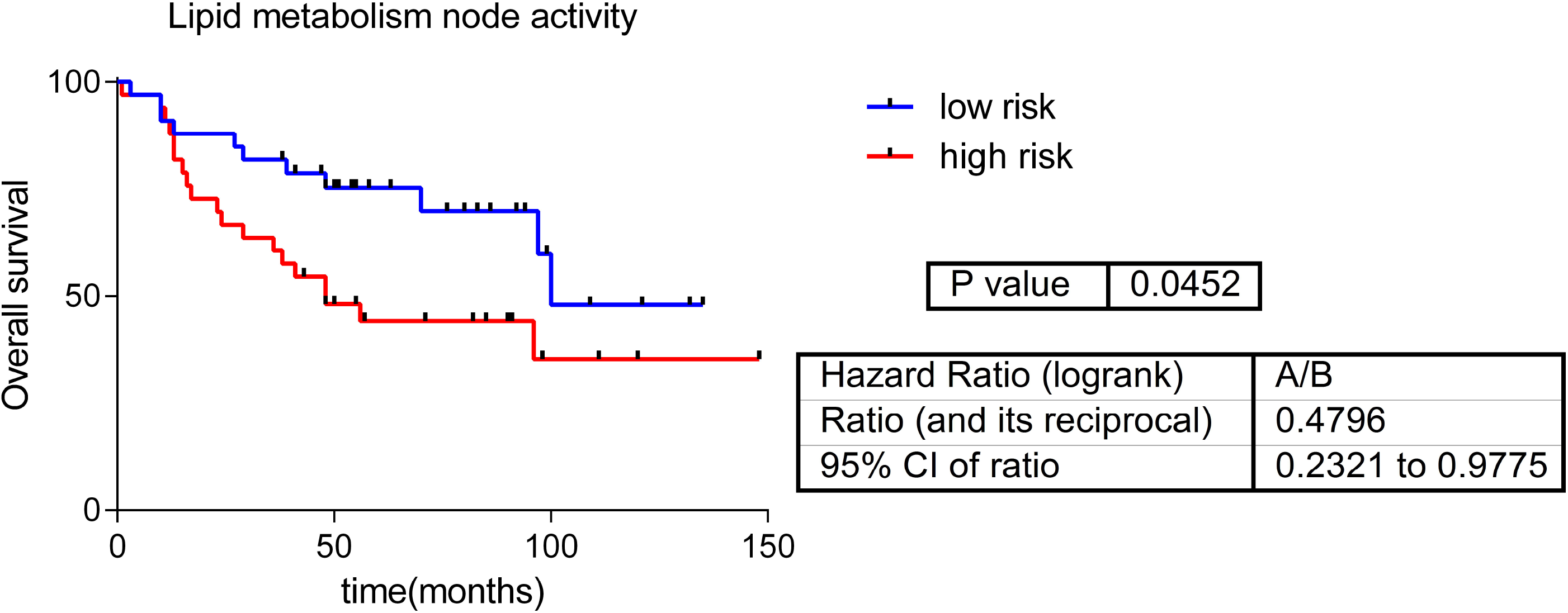
Predictor based on lipid metabolism node activity.

### Analyses combining gene expression with metabolomics data

On the other hand, a network combining metabolomics and gene expression data was built. Due to the differences between both kinds of data, most of the metabolites were grouped together. However, some metabolites were integrated into gene branches (Fig 4).

**Fig 4:**
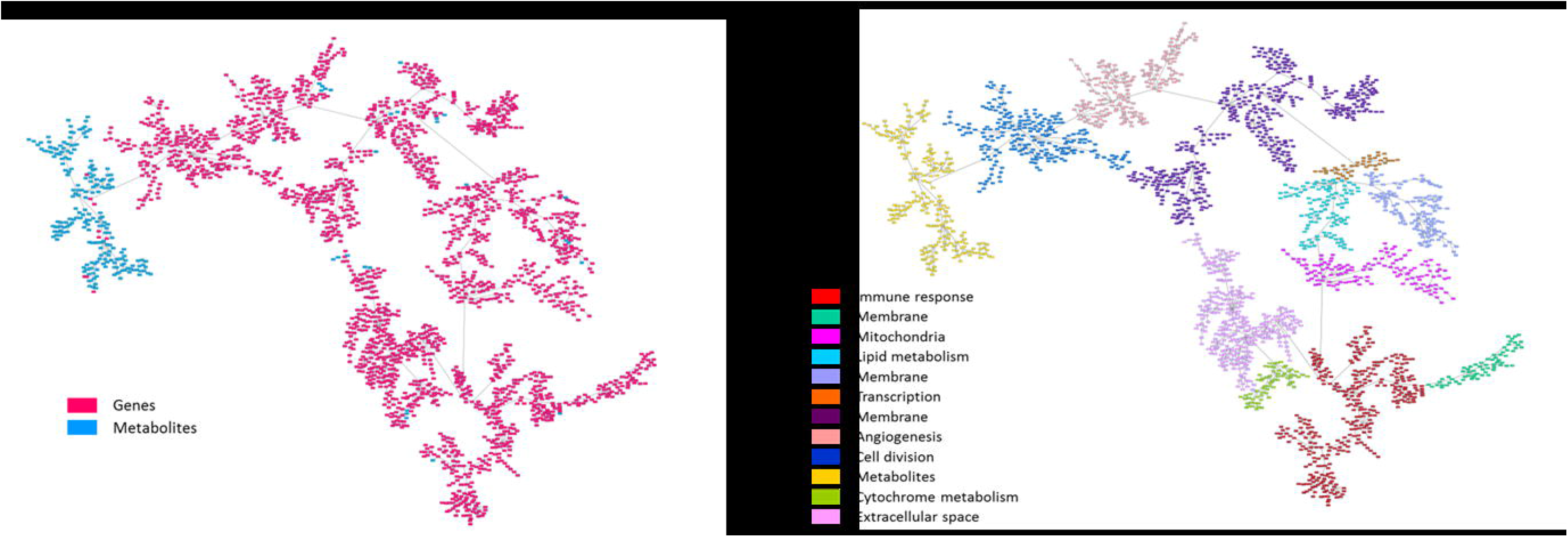
A. Network associating genes (red) and metabolites (blue). B. Metabolite and gene network functionally characterized.

This combined network was then functionally characterized based on the majority function of the genes contained in each branch. The resulting network had eleven functional nodes and a twelfth branch that include the majority of the metabolites (Fig 4).

Once the main functions were assigned, a literature review was performed to study the relationship between metabolites included in the gene functional nodes and the main function of each functional node. We found out that a relationship with functional nodes had been previously described for 4 of 20 metabolites: succinate, cytidine, histamine and 1,2-propanediol. The relationships between metabolites and their node function are shown in Table 2.

**Table 2:** Previously described relationships between metabolites included in gene nodes and the function of these nodes.

| Metabolite | Node | Described relationship | Reference |
| --- | --- | --- | --- |
| Succinate | Immune response | Increases immune response, induces IL-1b production, promotes adaptive immune response. | PMID: 28109906 |
| Cytidine | Immune response | 5-aza-2'-deoxycytidine potentiates antitumor immune response, role in innate immune response. | PMID: 23865062, PMID: 24559534 |
| Histamine | Angiogenesis | Histamine promotes angiogenesis by enhancing VEGF production. | PMID: 23225320 |
| 1,2-propanediol (prev.X-4796) | Angiogenesis | Modulates the immune system through S1P, which promotes angiogenesis and proliferation. 14C-sulfoquinovosyl acylpropanediol is an antiangiogenic drug. | PMID: 21632869, PMID: 29543539 |

### Flux Balance Analysis and flux activities

FBA and flux activities were calculated as previously described (Trilla-Fuertes et al, 2018). Briefly, gene expression data from 67 tumor samples were used, GPR rules were solved and the normalized values were introduced into the metabolic model using modified E-flux algorithm (Colijn et al, 2009; Gámez-Pozo et al, 2017). Finally, FBA was calculated using a biomass reaction representative of tumor growth. No significant differences were found in the tumor growth rate between ER+ and ER-tumors (S2 Fig).

Flux activities showed significant differences between ER+ and ER-in glycerophospholipid metabolism, phosphatidyl inositol metabolism, urea cycle, propanoate metabolism, pyrimidine catabolism and reactive oxygen species (ROS) detoxification (S3 Fig).

In addition, the combination of glutamate metabolism (the pathway that includes the glutamine) and alanine and aspartate metabolism flux activities showed prognostic value in this cohort (p-value = 0.024, HR = 0.41, 50:50%) (Fig 5). A multivariate analysis showed that this flux activity-based predictor provides prognostic information independently from clinical data (S3 Table).

**Fig 5:**
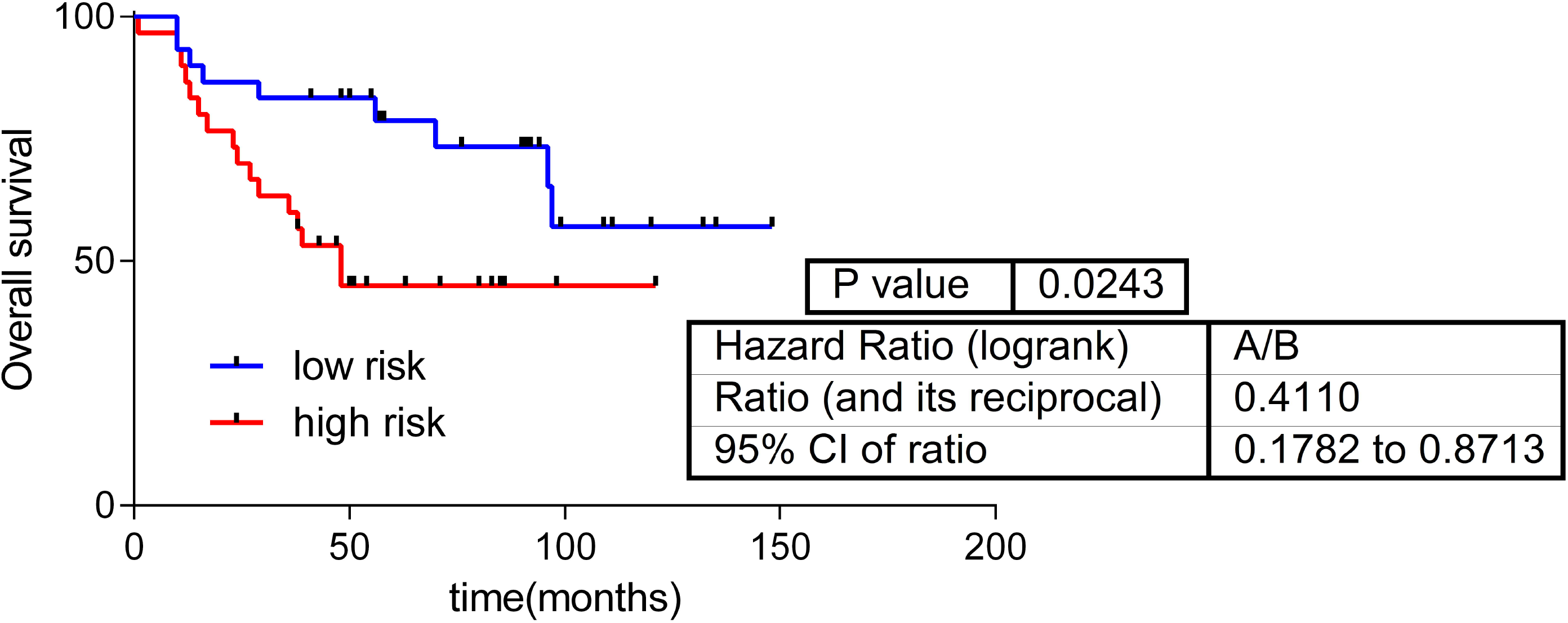
OS predictor based on glutamate metabolism and alanine and aspartate metabolism flux activities.

### PGM analysis combining flux activities with metabolomics data

Flux activities were calculated for each metabolic pathway defined in the Recon2. Then, using flux activities and metabolomics data, a new PGM was built to study association between both types of data. Interestingly, both types of data appeared mixed in the network; with the peculiarity that flux activities appeared usually at the periphery of the network (Fig 6).

**Fig 6:**
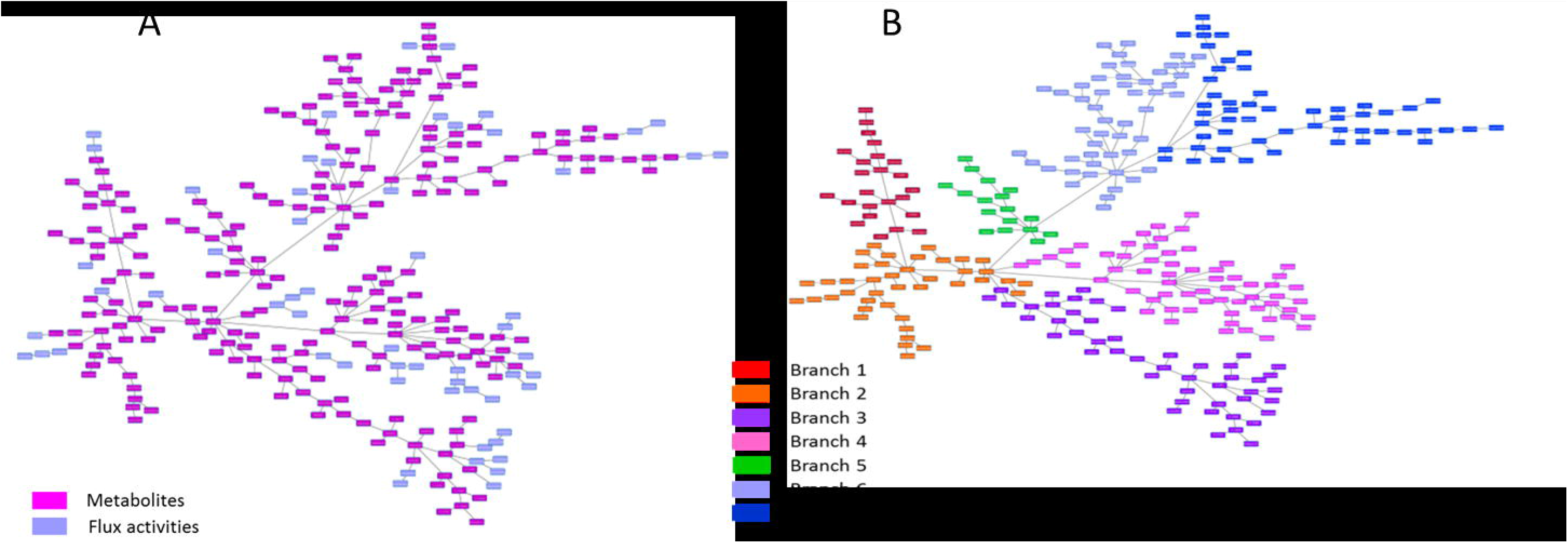
Probabilistic graphical model combining flux activities and metabolomics data. A. Network combining flux activities (purple) and metabolite (pink) expression data. B. Division in branches of the network formed by flux activities and metabolomics data.

The resulting network was split into several branches to study the relationship of the metabolites with the flux activities included in each branch (Fig 6). Coherence between both types of data was shown by the PGM, associating flux activities and metabolites related to these flux activities in the same branch. For instance, branch 1 includes glycolysis flux activity and three metabolites previously related to glycolysis (S4 Table). Regarding vitamins and cofactors, it was not possible make comparisons because the IMPaLA label for this category is “Vitamin and co-factor metabolism” and Recon2 labels differentiate between the various vitamins, labeling them as “Vitamin B6 metabolism”, “Vitamin A metabolism”, etc.

### Cell viability assays using drugs targeting glutamine metabolism

As the computational analyses pointed out the relevance of glutamine and its metabolism in breast cancer, cell viability assays employing two drugs targeting glutaminolysis (aminooxyacetic acid [AOA], and L-Glutamic acid γ-(p-nitroanilide) hydrochloride [GPNA]) were performed. Dose-response curves of these two drugs confirm that targeting glutamine metabolism affected cell viability (Figure 7).

**Figure 7:**
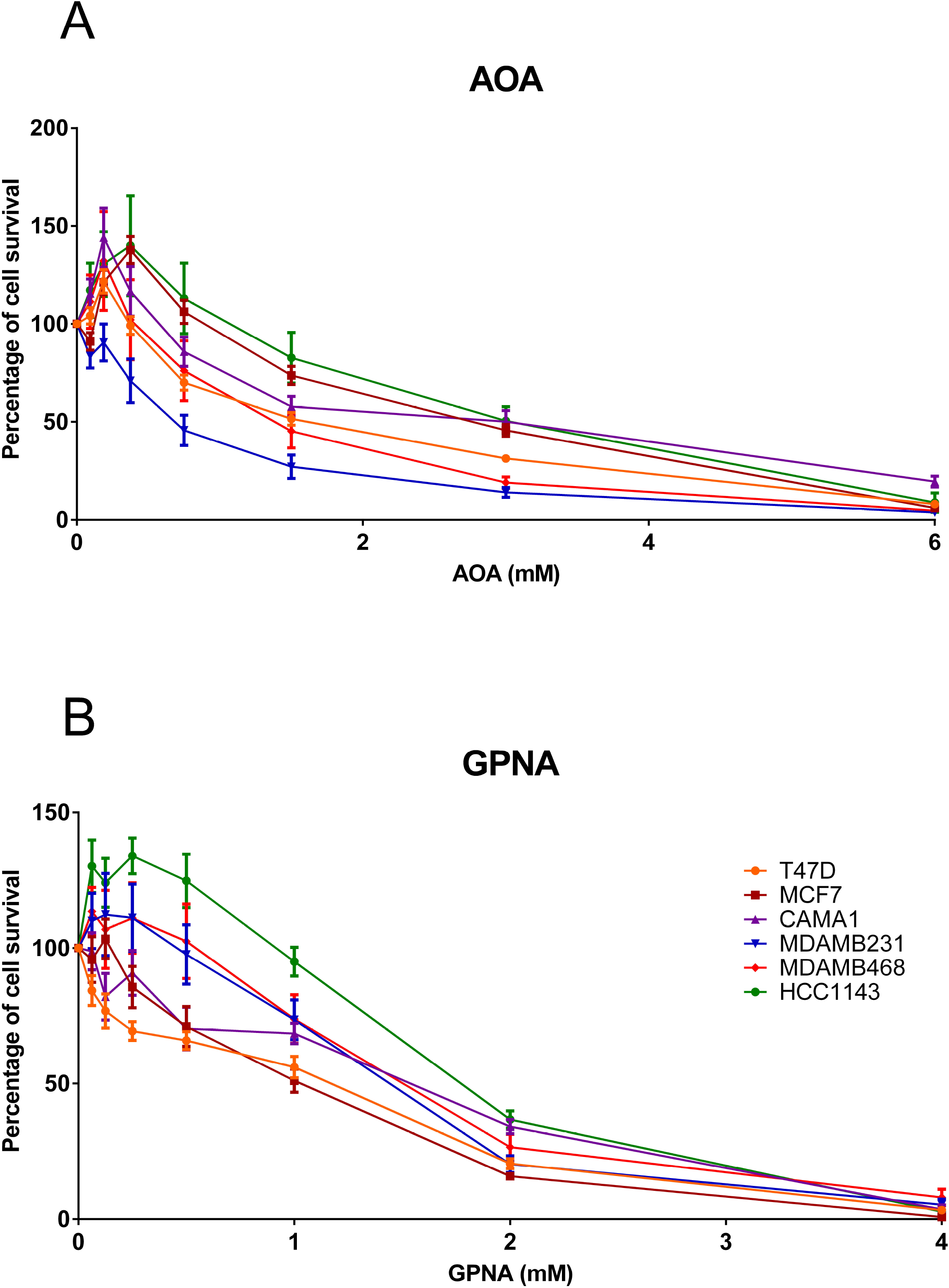
Dose-response curves using two drugs targeting glutamine metabolism in breast cancer cell lines. A. Dose-response curve for AOA (0-6 Mm). B. Dose-response curve for GPNA (0-4 Mm).

In AOA-treated cell lines, the IC_50_ did not show any differential response between breast cancer subtypes. However, in GPNA-treated cells, IC_50_ for ER+ cell lines were lower than IC_50_ for TNBC cells (Table 3).

**Table 3:** IC_50_ calculated for each drug in each breast cancer cell line.

| Cell Line | Subtype | AOA<br>IC<sub>50</sub> (mM) | GPNA<br>IC<sub>50</sub> (mM) |
| --- | --- | --- | --- |
| T47D | ER+ | 2.05 | 0.48 |
| MCF7 | ER+ | 3.89 | 0.69 |
| CAMA1 | ER+ | 2.90 | 1.10 |
| MDAMB231 | TNBC | 0.64 | 1.73 |
| MDAMB468 | TNBC | 2.29 | 2.50 |
| HCC1143 | TNBC | 4.22 | 2.59 |

## Discussion

Metabolomics is attracting considerable interest as a technique for finding new biomarkers in cancer. In this study, a new analytical workflow for the management and study of metabolomics data was proposed. This workflow allowed global metabolic characterization, beyond analyses based on unique metabolites. On the other hand, this workflow pointed out the relevance of glutamine metabolism in breast cancer, a hypothesis that was confirmed by cellular experiments.

Genomics and metabolomics data from Terunuma et al. have previously been used by The Cancer Genome Atlas Consortium to correlate gene expression data with metabolomics data (Peng et al, 2018; Terunuma et al, 2014). Based on this dataset, we applied PGMs for the first time in metabolomics data from tumor samples and also in metabolomics data combined with gene expression data and flux activities, with the aim of confirming known associations and finding new ones.

First, we evaluated whether metabolomics data were related to OS in patients with breast cancer. An OS predictive signature was built that included the expression values of glutamine, deoxycarnitine, butyrylcarnitine, glycerophosphorylcholine and 2-hydroxypalmitate (DeBerardinis et al, 2007). The first four metabolites have been previously related to survival in breast cancer (Bhowmik et al, 2015; Cao et al, 2012). However, to our knowledge, this is the first report associating 2-hydroxypalmitate with cancer survival. Additionally, in the previous study by Terunuma et al., 2-hydroxyglutarate was associated with poor prognosis in patients with breast cancer (Terunuma et al, 2014). 2-hydroxyglutarate is a glutamine intermediate in the tricarboxylic acid cycle, involved in the conversion of glutamine into lactate, a process known as glutaminolysis (DeBerardinis et al, 2007). The negative sign in the predictor of glutamine points a protective effect (the more glutamine the better prognosis). An increased presence of glutamine could indicate that it has not been introduced into the Krebs cycle and transformed into lactate, a fact associated with a more aggressive phenotype and a worst prognosis.

A metabolite network using metabolomics data was built using PGMs. PGMs are based on expression data, or quantification data in the case of metabolomics, and they do not need any additional information. The output of this analysis is a network that reflects the correlations between genes, proteins or metabolites. On the other hand, IMPaLA assigned a dominant metabolic function to each resulting node. In previous studies, we demonstrated that PGMs are useful for functionally characterizing gene or protein networks (de Velasco et al, 2017; Gámez-Pozo et al, 2015; Gámez-Pozo et al, 2017). However, to our knowledge, this is the first time a PGM has been applied to metabolomics data from tumor samples. Just as observed in genes or proteins, metabolites are grouped into metabolic pathways, allowing the characterization of differences in metabolic pathways between ER+ and ER- tumors. For example, both lipid metabolism and purine metabolism node activities were higher in ER- tumors. Although there has not been described a relationship between lipids and breast cancer subtypes, it was described that ER-tumors usually overexpress genes related to lipid metabolism (Wang et al, 2017). Moreover, the activity of the lipid metabolism node had prognostic value. No relationship between purine metabolism and breast cancer has previously been defined.

On the other hand, the network combining gene expression data and metabolomics data grouped most of the metabolites into an isolated branch. Yet, some metabolites were included into gene branches. We found that four out of twenty metabolites showed a previously reported relationship with the main function of the gene functional node in which they were included. Succinate and cytidine were located in the immune response functional node. Succinate acts as an inflammation activation signal, inducing IL-1β cytokine production through hypoxia-inducible factor 1 (Tannahill et al, 2013). In addition, succinate increases dendritic cell capability to act as antigen-presenting cells, prompting an adaptive immune response (Jiang & Yan, 2017). Regarding cytidine, Wachowska et al. described that 5-aza-2’-deoxycytidine modulates the levels of major histocompatibility complex class I molecules in tumor cells, induces P1A antigen and has immunomodulatory activity when combined with photodynamic therapy (Wachowska et al, 2014).

Both histamine and 1,2-propanediol appeared to be related to the angiogenesis functional node. Histamine is known to promote angiogenesis through vascular epithelial growth factor (Lu et al, 2013). On the other hand, sulfoquinovosyl acylpropanediol, an 1,2-propanediol derivate, inhibits angiogenesis in murine models with pulmonary carcinoma (Ruike et al, 2018).

The remaining sixteen metabolites require an in-depth study to establish associations with their respective functional nodes. These results support the potential of PGMs as a tool to generate hypotheses without the need of *a priori* knowledge.

FBA was used to model metabolism using gene expression data. Although FBA-predicted biomass did not show significant differences between ER+ and ER-tumors, differences in flux activities were shown between both subtypes. Some of these activities were also related to prognosis, such as “Glutamine metabolism”, which agrees with the results obtained from the metabolomics data, including glutamine in the metabolite signature capable of predicting OS. These results highlighted the relevance of glutamine metabolism in breast cancer, suggesting the utility of drugs targeting this pathway such as GPNA, which it has already been described as affecting proliferation in lung cancer cells (Hassanein et al, 2013). Strikingly, cell viability experiments using GPNA and AOA, two drugs targeting glutamine metabolism, showed a decreased in cell viability, confirming the relevant role of this process in breast cancer. Despite the highest levels of glutamine-related enzymes described in TNBC and HER2 tumors comparing with luminal tumors, dose-response curves did not show any differential response between ER+ and TNBC breast cancer cells in the case of AOA (Cao et al, 2014; Kanaan et al, 2014). In addition, ER+ cells seem to be more sensible to GPNA. On the other hand, AOA has been successfully tested in ER+ and ER-breast cancer xenograft models (Korangath et al, 2015).

With the aim of associating metabolomics and FBA results, flux activities and metabolomics data were combined to form a new network. As opposed to gene and metabolite data, metabolomics data and flux activities combined well in the network. Interestingly, flux activities are dead-end nodes, perhaps due to the fact that they are by definition a final summary of each pathway. IMPaLA assigned a main metabolic pathway to resulting branches; thus, it was possible to know how many metabolites were related to flux activity in each branch. In most cases with available information, there was coherence between metabolites included in the branch and its flux activity. This validates FBA and flux activities, both based on gene expression, as a method of simulating metabolism.

Our study has some limitations. The limited number of samples leads us to consider the results as preliminary, and validation in an independent cohort is needed. Additionally, our results are difficult to place in the current clinical landscape, given that tumors in the original series had not been assessed for HER2 expression. On the other hand, evolving techniques currently allow the detection of more metabolites, which would permit a more thorough analysis.

Metabolomics is postulated as a booming technique for the biomarker search in cancer. Additionally, PGMs reveal their utility in the analysis of metabolomics data from a functional point of view, not only metabolomics data alone, but also in combination with flux or gene expression data. Therefore, PGM is postulated as a method to propose new hypotheses in the metabolomics field. We also found that it is possible to associate metabolomics data with clinical outcomes and to build prognostic signatures based on metabolomics data. Finally, these computational analyses suggested a main role of glutamine metabolism in breast cancer, a fact that was experimentally validated.

## Materials and methods

### Patients included in the study

Metabolomics and gene expression data from 67 fresh tumor tissue samples originally analyzed by Terunuma et al. (Terunuma et al, 2014) were included in this study.

### Preprocessing of gene expression and metabolomics data

Metabolomics data contains information about 536 metabolites. Log2 was calculated. As quality criteria, data were filtered to include detectable measurements in at least 75% of the samples. Missing values were imputed to a normal distribution using Perseus software (Tyanova et al, 2016). After quality control, 237 metabolites were considered for subsequent analyses.

In terms of gene expression data, the 2,000 most variable genes, i.e., those genes with the highest standard deviation, were chosen to build the PGM. This data was from an Affymetrix array and they are available in Gene Expression Omnibus Database under the identifier GSE37751.

### Probabilistic graphical models and gene ontology analyses

As previously described (de Velasco et al, 2017; Gámez-Pozo et al, 2015; Gámez-Pozo et al, 2017; Trilla-Fuertes et al, 2018), PGMs compatible with high dimensional data were used, using correlation as associative criteria. PGMs were built using metabolomics, gene expression or flux activity data without any a priori information. The *grapHD* package (Abreu et al, 2010) and R v3.2.5 were employed to build the PGMs. The management of the resulting network was done using Cytoscape software (Shannon et al, 2003). The resulting networks were divided into branches and ontology analyses were done to assign a majority function/metabolic pathway to each branch, defining in this way different functional nodes in the networks. In the case of genes, gene ontology analyses were performed using the DAVID web tool with “homo sapiens” as background and GOTERM, KEGG and Biocarta selected as categories (Huang et al, 2009). In the case of metabolites, the Integrated Molecular Pathway Level Analysis (IMPaLA) web tool was used to assign a main metabolic pathway to each branch (Cavill et al, 2011).

Once the functional structure was defined, functional node activities were calculated in order to make comparisons between groups, as previously described (de Velasco et al, 2017; Gámez-Pozo et al, 2015; Gámez-Pozo et al, 2017; Trilla-Fuertes et al, 2018). Briefly, each functional node activity was calculated as the mean of the expression/quantity of genes/metabolites of each node that are related to the main node function/metabolic pathway.

### Flux Balance Analysis and flux activities

FBA is a method that allows the estimation of tumor growth rate. FBA was performed using the library COBRA Toolbox v2.0 (Schellenberger et al, 2011) available for MATLAB. FBA was calculated using the whole human metabolic reconstruction Recon2 (Thiele et al, 2013). This metabolic reconstruction includes 2,191 genes collected into the Gene Protein Reaction rules (GPRs), 5,063 metabolites and 7,440 reactions. GPRs represent the relationships between genes and metabolic reactions and they are included into the model as Boolean expressions. GPRs were solved as described in previous studies (Gámez-Pozo et al, 2017; Trilla-Fuertes et al, 2018), using a modification of the Barker *et al*. algorithm (Barker et al, 2015), which were incorporated into the model by a modified E-flux method (Colijn et al, 2009; Trilla-Fuertes et al, 2018). Briefly, the “OR” operators were solved as the sum and the “AND” operators were solved as the minimum. Then, the GPR data were normalized using the Max-min function and introduced into the model as the reaction bounds. As the objective function, the biomass reaction proposed in the Recon2 was used as representative of tumor growth. This biomass reaction was based on experimental measurements of leukemia cells. The 7,440 reactions are grouped into 101 metabolic pathways.

Flux activities were previously proposed as a measurement to compare differences at the metabolic pathway level (Trilla-Fuertes et al, 2018). Briefly, they were calculated as the sum of the fluxes of the reactions included in each pathway defined in Recon2.

### Statistical analyses

The statistical analyses were performed with GraphPad Prism v6. Predictor signatures were built with the BRB Array Tool from Dr. Richard Simon’s team (Simon, 2005). All p-values are two-sided and are considered statistically significant under 0.05.

### Cell culture and reagents

Breast cancer cell lines (MCF7, T47D and CAMA1 [ER+], and MDAB231, MDAMB468 and HCC1143 [TNBC]) were cultured in RPMI-1640 medium with phenol red, supplemented with 10% heat-inactivated fetal bovine serum, 100 mg/mL penicillin and 100 mg/mL streptomycin. Cell lines were cultured at 37°C in a humidified atmosphere with 5% (v/v) CO2 in the air. The MCF7, T47D and MDA-MB-231 cell lines were kindly provided by Dr. Nuria Vilaboa (La Paz University Hospital, previously obtained from ATCC in January 2014). The MDAMB468, CAMA1 and HCC1143 cell lines were obtained from ATCC (July 2014). Cell lines were routinely monitored and authenticated, tested for Mycoplasma and frozen, and passaged for fewer than 6 months before experiments. The AOA (Sigma Aldrich C13408) and GPNA (Sigma Aldrich G6133) were obtained from Sigma-Aldrich (St. Louis, MO, USA).

### Cell viability assays

Dose-response curves were designed for AOA and GPNA. As the preparation of GPNA needs an acid medium, HEPES (50mM) was added to buffer the medium. About 5,000 cells were seeded in each well in 96-well plates and after 24 hours, drugs were added. After an incubation of 72 hours, cell viability was determined using CellTiter 96 AQueous One Solution Cell Proliferation Assay (Promega) kit and absorbance was quantified on a microplate reader (TECAN). As a control untreated cells were used and all the experiments were performed by triplicate.

## Acknowledgements

This study was supported by Instituto de Salud Carlos III, Spanish Economy and Competitiveness Ministry, Spain and co-funded by the FEDER program, “Una forma de hacer Europa” (PI15/01310). LT-F is supported by the Spanish Economy and Competitiveness Ministry (DI-15-07614). GP-V and EL-C are supported by Consejería de Educación, Juventud y Deporte of Comunidad de Madrid (IND2017/BMD7783). The funders had no role in the study design, data collection and analysis, decision to publish or preparation of the manuscript.

## Author contributions

All the authors have directly participated in the preparation of this manuscript and have approved the final version submitted. JMA, MD-A, HN and PM contributed the directed graphical models. LT-F, AG-P, G-PV, and AZ-M performed the statistical analyses, the graphical model interpretation and the ontology analyses. LT-F, AG-P, JAFV, PZ and EE conceived of the study and participated in its design and interpretation. MD-A and LT-F performed the Flux Balance Analysis. EL-C did the cell viability experiments. LT-F drafted the manuscript. AG-P, JAFV, and EE supported the manuscript drafting. AG-P and JAFV coordinated the study.

## Data Availability Section

Not applicable.

## Supporting information

**S1 Table: Multivariate Cox regression model comparing OS predictor based on metabolomics data.** T = tumor stage, N = lymph node status, G = tumor grade.

**S2 Table: Multivariate Cox analysis comparing predictor based on node activity of lipid metabolism.** T = tumor stage, N = lymph node status, G = tumor grade.

**S3 Table: Multivariate Cox regression comparing predictor based on flux activities.** T = tumor stage, N = lymph node status, G = tumor grade.

**S4 Table: Metabolites associated with flux activity of each network branch.**

**S1 Fig: Node activities from the metabolic network.**

**S2 Fig: Tumor growth rate predicted using FBA for ER+ and ER- tumors.**

**S3 Fig: Flux activities were significantly different between ER+ and ER-.**

