## Supplementary material for "Computational metabolomics hints at the relevance of glutamine metabolism in breast cancer": Supplemmentary Table 1

|  | Parametric p-value | FDR | Hazard Ratio | UniqueID |
| --- | --- | --- | --- | --- |
| 1 | <0.001 | 0.159 | 0.3 | glutamine |
| 2 | 0.002 | 0.179 | 2.0 | 2-hydroxypalmitate |
| 3 | 0.004 | 0.212 | 1.5 | deoxycarnitine |
| 4 | 0.005 | 0.212 | 1.3 | glycerol 3-phosphate (G3P) |
| 5 | 0.006 | 0.212 | 1.5 | butyrylcarnitine |
| 6 | 0.007 | 0.212 | 1.7 | carnitine |
| 7 | 0.007 | 0.212 | 1.7 | inosine |
| 8 | 0.007 | 0.212 | 1.6 | 3-dehydrocarnitine* |
| 9 | 0.008 | 0.212 | 1.3 | palmitoylcarnitine |
| 10 | 0.010 | 0.212 | 1.3 | S-adenosylhomocysteine (SAH) |
| 11 | 0.011 | 0.212 | 1.8 | creatine |
| 12 | 0.012 | 0.212 | 1.3 | 2-linoleoylglycerophosphocholine* |
| 13 | 0.012 | 0.212 | 1.6 | guanosine |
| 14 | 0.013 | 0.212 | 1.7 | X - 11787 |
| 15 | 0.014 | 0.212 | 1.2 | stachydrine |
| 16 | 0.015 | 0.212 | 1.4 | X - 12855 |
| 17 | 0.015 | 0.212 | 1.8 | 2-hydroxystearate |
| 18 | 0.019 | 0.249 | 1.4 | hexanoylcarnitine |
| 19 | 0.022 | 0.278 | 1.4 | 22:6n3);docosahexaenoate (DHA |
| 20 | 0.024 | 0.283 | 1.2 | N-acetylaspartate (NAA) |
| 21 | 0.026 | 0.294 | 1.2 | 2-oleoylglycerophosphocholine* |
| 22 | 0.029 | 0.311 | 1.2 | oleoylcarnitine |
| 23 | 0.030 | 0.311 | 1.3 | ribose |
| 24 | 0.032 | 0.311 | 1.4 | glycerophosphorylcholine (GPC) |
| 25 | 0.033 | 0.311 | 1.2 | X - 12627 |
| 26 | 0.035 | 0.322 | 1.5 | succinate |
| 27 | 0.039 | 0.345 | 1.3 | X - 3094 |
| 28 | 0.046 | 0.378 | 1.3 | propionylcarnitine |
| 29 | 0.048 | 0.378 | 1.2 | 2-arachidonoylglycerophosphocholine* |

Sup Table 1: Metabolites associated with overall survival in Terunuma et al. cohort (p<0.05).
