## Supplementary figures and images for "Computational metabolomics hints at the relevance of glutamine metabolism in breast cancer"

### Supplemmentary Figures

Sup Fig 1

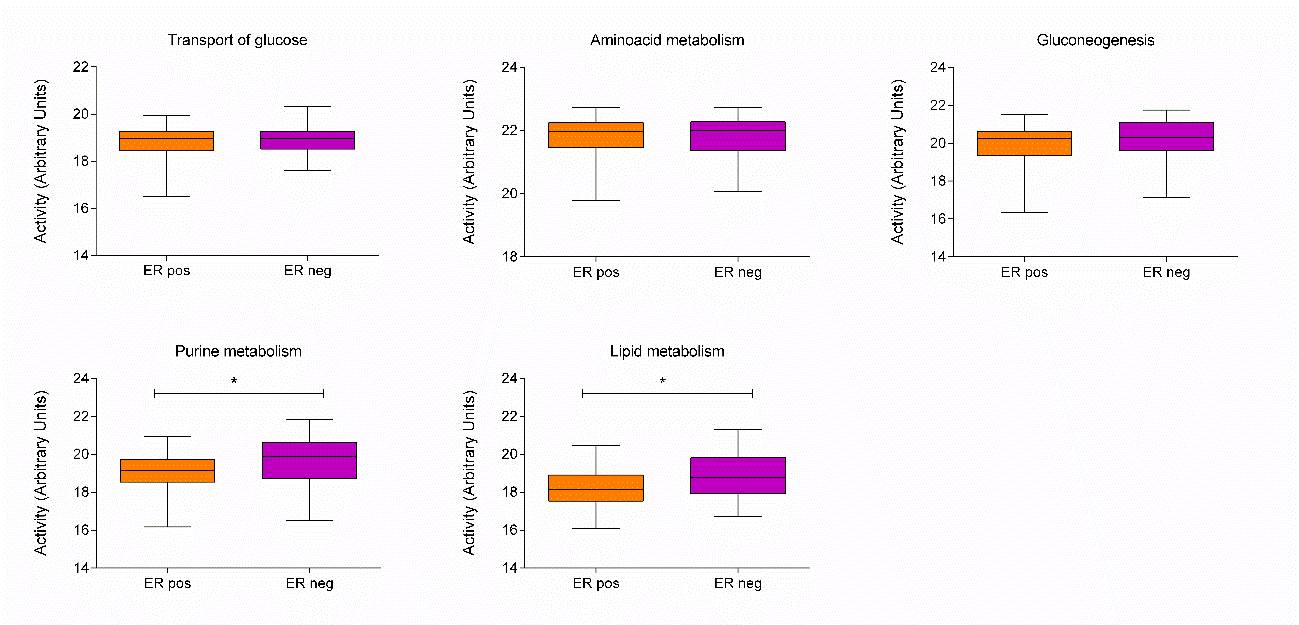

Sup Fig 2

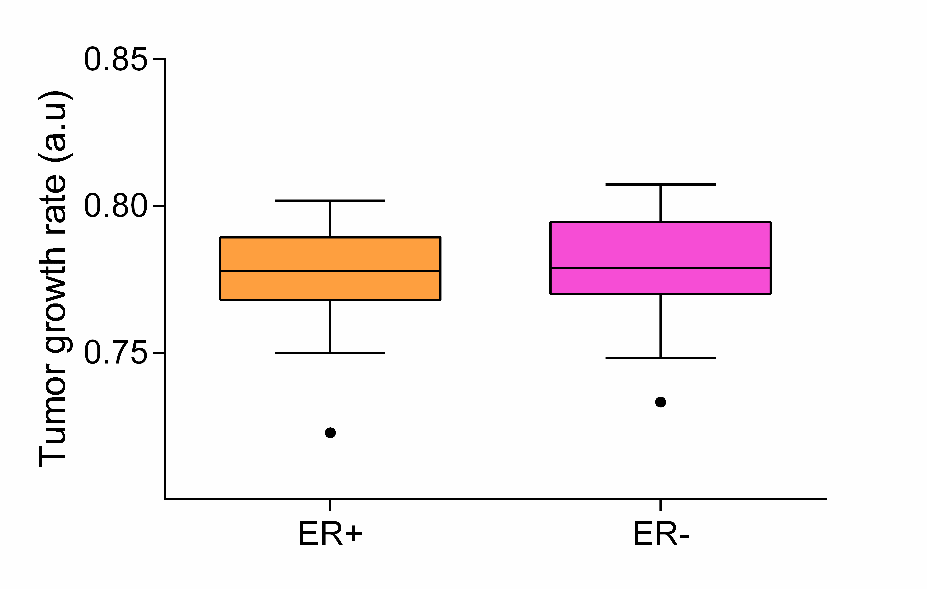

Sup Fig 3

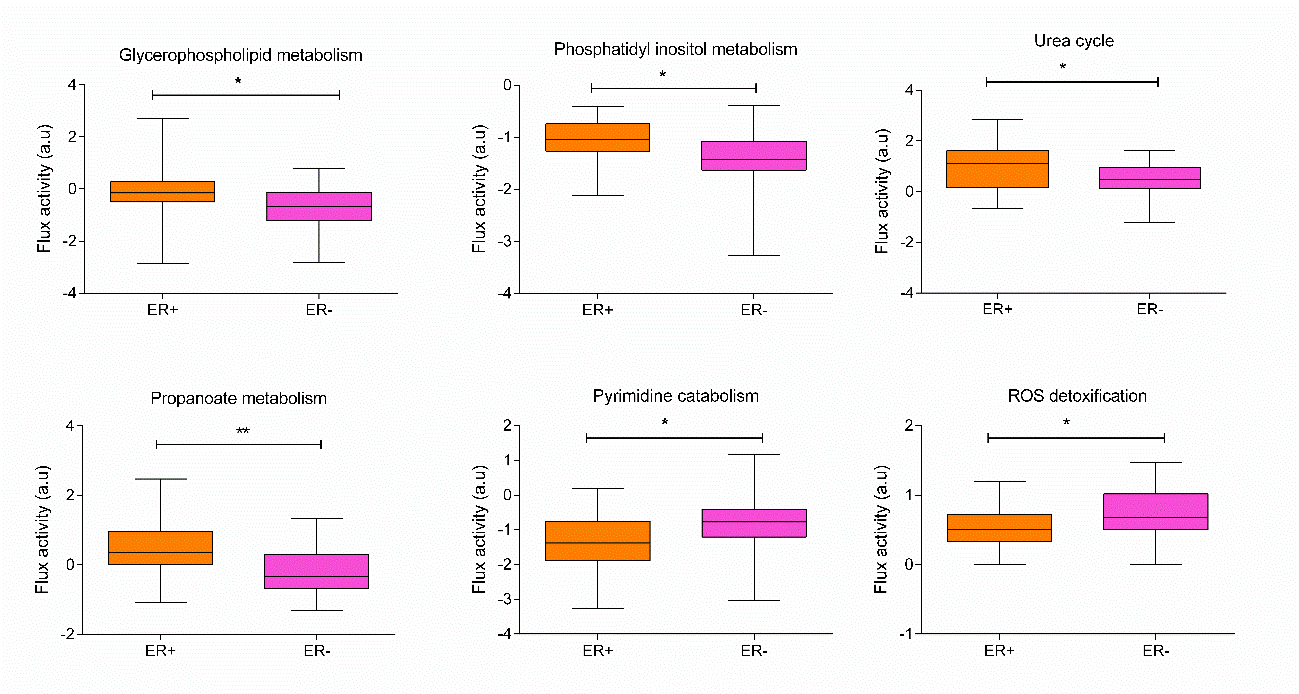
